# Equine Facial Action Coding System for determination of pain-related facial responses in videos of horses

**DOI:** 10.1101/2020.03.31.018374

**Authors:** Maheen Rashid, Alina Silventoinen, Karina B. Gleerup, Pia H. Andersen

## Abstract

During the last decade, pain scales including facial expressions as indicators of pain have been developed for horses, mostly relying on direct observations or inspection of images. Despite differences in the research conditions and methodology the different scales focus on the same regions of the face, corresponding to moveable facial muscles related to the ears, eyes, nostrils, lips and chin. However, a detailed comparison of the facial activities occurring during pain is not possible. We used a Facial Action Coding System modified for horses to code and analyse video recordings from an earlier study of acute short-term experimental pain and from clinical cases with and without pain. We demonstrated for the first time EquiFACS based changes to pain in video of horses, using traditional statistical methods based on frequency, and novel analyses based on sliding observation windows and co-occurrence of facial actions. The most prominent differences of the experimental horses were related to the lower face actions *chin raiser* and *nostril dilator*, while less prominent, but significantly more frequent actions were related to the eye region, *inner brow raiser* (AU101), *increased eye white* (AD1), *half blink* (AU47), and *ear rotator* (EAD104). *Ears forward* (EAD101) and *eye blink* (AU145) were not associated to pain. Based on this we selected the two lower face actions for analysis of the clinical videos, and found that their co-occurrence within a window of 10 to 15 second gave 100% positive predictive values, as compared to the rating from three expert pain raters. Using our developed co-occurrence analyses we were surprised to detect that the chance of identifying three or more of the facial actions related to pain in 0.04 second sequence, corresponding to one frame, was below 3%, indicating that use of randomly selected images for pain scoring may be a very insensitive method.

## 2 Introduction

Pain is a sign of disease, and early recognition of pain may improve welfare and treatment of otherwise disabling diseases in horses. Pain is an unpleasant sensory and emotional experience associated with actual or potential tissue damage, as currently defined by the International Association for the Study of Pain (IASP) [1].While self-reporting is the gold standard for assessment of pain in verbal humans [2], there are no affective measures available for the aversive components of pain in non-verbal mammals, including the horse [3]. The IASP definition adds that “the inability to communicate verbally does not negate the possibility that an individual is experiencing pain”, referring to adults unable to communicate, neonates and infants, as well and animals. This has brought attention to communication conveyed by non-verbal behaviours, such as bodily behavior, visible physiological activity such as muscle tremor and facial expressions. During the last decades a plethora of pain scales based on pain-related bodily behavior has been developed for horses [4–9], while research in facial expressions as indicators of pain in horses is a more recent contribution [10, 11]. In the Gleerup study [11], pain was induced in otherwise healthy and trained horses using short-term acute pain induction models, whereas the horses in the Dalla Costa study [10] experienced postoperative pain from castration. Despite many differences in the conditions and methodology, the two studies identified and described facial activity in the same regions of the face, corresponding to moveable facial muscles related to the ears, eyes, nostrils, lips and chin. However, differences were also present, for example regarding orbital tightening, which was observed only in some studies. Dalla Costa et al. [12] later identified a classifier that could estimate the pain status of the animal based on the facial activities coded from still images in the Horse Grimace Scale and thus confirming that the individual scores were related to the pain state of the horse.

Due to differences in both experimental approaches and descriptions of the facial activities observed, a detailed comparison of the facial activities occurring during pain have not been possible.

In humans, the Facial Action Coding System (FACS) provides a recognized method for identifying and recording facial expressions based on the visible movement of the underlying facial muscles in humans [13]. The coding requires training, and reliable coding can be expected from certified coders.

Recently, Wathan et al. [14], on the basis of this FACS methodology, developed the Equine Facial Action Coding System (EquiFACS) for horses. EquiFACS exhaustively describes all observable equine facial behavior using AUs, Action Descriptors (ADs) and Ear Action Descriptors (EADs). FACS coding relies on detailed video observation of facial muscle movement as well as changes in facial morphology (e.g., the position of the eyebrows, size/shape of the mouth, lips, or eyelids, the appearance of various furrows, creases, bulges of the skin) to determine which AU(s) occurred. Inter-observer agreement is good-to-excellent for spontaneously generated facial behavior in 90% of the action units in humans [15] scored by trained and certified FACS readers.

The work by Wathan [14] argues that facial movements can only be coded accurately from video sequences, and information from FACS annotations ideally contains precise information about times of onset, peak and offset of the individual AUs. The FACS systems exhaustively code all facial activity observed, not only what is thought to be pain-related. Any interpretations of the emotional meaning of the observed AUs occur post-coding, so the coding system itself is entirely atheoretical.

Pain-related facial responses in horses has never been described using this finely-grained and objective coding system. While the methodology exists for the coding of horse facial activity, no methods exist for the interpretation of the results.

Therefore, the aim of this study was to code facial expressions of horses before and during acute experimental pain, and to develop and test statistical approaches that define pain-related facial movements in EquiFACS.

## 3 Horse Pain Dataset

### 3.1 Experimental Pain Data

We used the videos recorded from an earlier study [11] including six healthy horses of different breeds, five mares and one gelding. Briefly, the horses were stabled at the research facility for at least ten days before the study, and were positively reinforced during this time to stand in the trial area while wearing only a neck collar. These conditions were designed to increase the horses comfort in trial settings, reducing the risk of external factors influencing the horse. Baseline recordings were obtained on the day of the experiment. Acute short term ischemic pain was induced by the application of a pneumatic blood pressure cuff placed on a forelimb for up to 10 minutes.

From the raw video footage, video clips of 30 seconds duration were selected from the base line period (first occasion where the profiled horse were within the frame for 30 secs) and first occasion that the profiled horse head were within the frame during maximum experience of pain.

### 3.2 Clinical Pain Data

The horses in the experimental data described above were habituated to the surroundings and filming conditions. Therefore, videos in this dataset are unlikely to contain signs of fear, stress or other emotional states commonly correlated with pain, and can be considered clean and reliable. While useful for developing a model for pain detection, the data is a restrictive and distant from the natural setting for pain evaluation.

We therefore collect additional data in a clinical setting. This data is used to evaluate our models’ ability to generalize to a less controlled setting. Twenty-one horses admitted to a horse hospital for treatment of both presumed painful diseases or non-painful conditions were filmed with owners’ consent for research purposes, but not for publication.

30 second clips from each video were EquiFACS annotated. Three veterinary experts assessed pain level based on their clinical experience as one of ‘Severe Pain’, ‘Moderate Pain’, and ‘No Pain’ for each horse. To obtain a single pain label, we used majority voting between raters. That is, if at least two of the three raters labeled a video as either ‘Moderate’ or ‘Severe’ pain, the video was labeled as ‘Pain’, else if was labeled as ‘No Pain’. This results in 7 pain and 14 no-pain videos.

### 3.3 Equine Facial Action Coding System

Equine Facial Action Coding System, as described by Wathan et al. [14] was used for a complete annotation. The system consists of 16 Action Units (AUs), and 11 Action descriptors (ADs), of which 4 are Ear Action Descriptors (EADs). While AUs represent the contraction of a particular muscle or muscle group, Action Descriptors describe a more general movement caused by either an undetermined muscular basis, or by deep muscles. *For simplicity all EquiFACS codes are named Action Units in the text*.

All films were coded in a blinded manner by a certified EquiFACS coder without knowledge of the study horses. The annotation program ELAN [16] was used to annotate start and stop of all Action Units (AU), Action Descriptors (ADs) and Ear Action Descriptors (EADs). As a result the annotated dataset contains not only the occurrence of different action units, but also the duration of time they are active and their overlap in time with other active action units.

## 4 Discovering Pain AUs

The EquiFACS dataset derived from the videos was used to identify the action units most useful for the identification of pain in a data-driven manner. Statistical patterns in AU occurrence were used to derive a set of action units that can differentiate between horses with pain and horses without pain. AUs associated with head and neck movement were excluded as they do not correspond with facial expressions.

We use a paired t-test for mean values to test significance. The number of times an AU occurs within an observation is used for the t-test.

### 4.1 Human FACS Interpretation (HFI) Method

While studies applying EquiFACS methods are few, research on *human* facial expressions of pain is mature and extensive. Kunz et al [17] presents a systematic review of studies on human facial expressions of pain and describe current approaches for the identification of action units associated with pain. The first criterion for defining an AU to be pain related is if it occurs frequently – forming more than 5% of total pain AU occurrences [18]. The second approach is to define an AU as pain related if it occurs more frequently during pain than during baseline [19]. Often both criteria are applied after each other [20], resulting in a set of AUs that are both frequent and distinct.

We use the two step approach in determining pain AUs - first AUs that form more than 5% of all pain AU occurrences are selected. From these the AUs that occur more frequently in pain rather than no-pain videos are determined as a the final pain related AUs.

### 4.2 Co-occurrence Method

While the method presented above (Section 4.1) is effective and simple also for horses, it does not take into consideration the temporal distribution of onset-offset of the various action units. The recognition of the facial expressions of pain constitute a selections of action units present at the same time or in close relation to each other. We therefore hypothesize that pain expressions are defined not by the occurrence of individual action units, but rather by multiple action units occurring together in the same period of time.

AUs that comprise a pain expression are likely to co-occur, i.e. occur together, in a pain state, and are likely to co-occur with a different set of AUs in a no-pain state. By comparing patterns of co-occurrence between pain and no-pain states, we can discover AUs most indicative of a pain expression.

To achieve this, we build a graph to capture the co-occurrence relationships between AUs. Each node represents an AU and edges between nodes are weighed by how often they occur together. We then inspect how edge weights change between pain and no-pain videos, and select the AUs that exhibit the largest change as pain AUs. All AUs that are active during a pre-defined slice of time (or Observation Window) are counted as co-occuring. This information is available since we record the start and end time of each AU’s activation (see Section 3.3).

More specifically, we build a ‘Co-occurrence Graph’ each for pain – *G*_*P*_ – and no-pain – *G*_*NP*_ –states. The graph is represented as a *N* × *N* adjacency matrix, where *N* is the total number of annotated action units, and value in row *i* and column *j* of the matrix represents the fraction of times AU *j* occurs in the same slice of time as action unit *i*. For example, if AU *j* occurs together with AU *i* in 5 time slices, and AU *i* occurs in 10 time slices in total the value in row *i*, column *j* – referred to as *G*^*i,j*^ – would be 5*/*10 = 0.5. The diagonal of this matrix is set to zero. The Co-occurrence Graph is directed, meaning that the value in *G*^*i,j*^ need not equal *G*^*j,i*^ since AU *i* and *j* can occur in a different total number of time slices.

Using fraction, or relative co-occurrence, rather than raw co-occurrence count to weigh each edge acts as a normalization procedure such that AUs that occur more frequently (such as blinking) do not have higher edge weights than AUs that occur less frequently. Edge values also become easily interpretable as they capture the co-occurrence rate of any two AUs relative to other co-occurring AUs, and are bounded between 0 and 1.

Following, we subtract the adjacency matrix of no-pain co-occurrence graph from the adjacency matrix of pain co-occurrence graph to get a ‘Difference Graph’, *G*_*D*_.

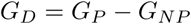

*G*_*D*_ captures changes in relative co-occurrence importance between pain and no-pain states. For example a difference value of +0.3 between AU *i* and *j* implies that AU *j* constitutes 30% more of all co-occurrences in pain than it did in no-pain for AU *i*, and has increased in relative co-occurrence importance. Note that since the pain and no-pain Co-occurrence Graphs are directed graphs, the Difference Graph is also directed.

AUs with the largest values in the Difference Graph are considered important for pain detection. Given an AU, we calculate its total change in relative co-occurrence importance for all its co-occurring AUs between pain and no-pain states. AUs that show more than median change are chosen as pain AUs.

More formally, let 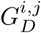 be the value in the *i*th row and *j*th column in the difference graph adjacency matrix. The importance of AU *i* to pain detection,*r*_*i*_, is then calculated by using the following formula that sums column *i*:

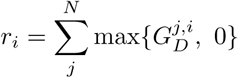

In other words, *r*_*i*_ sums the change in relative co-occurrence AUs that co-occur with AU *i* experience. Using column wise summation helps highlight AUs that influence the relative co-occurrence of other AUs. A row wise summation, on the other hand, would disproportionately highlight AUs that only exhibit large changes in relative co-occurrence because they occur once or close to once in the entire dataset. By ignoring decreases, or negative values, in the summation, we avoid AUs that are negatively correlated with pain.

#### 4.2.1 Conjoined Pain AUs

The AU selection method described above simply selects AUs that exhibit a large change in relative co-occurrence. The selected AUs need not occur together in the same time slice. However, AUs that comprise a pain expression must occur together in the same time slice. They should also occur more frequently in pain rather than no-pain states. We refer to these as conjoined AUs.

This equates to finding a cluster of AUs in the Difference Graph that are firstly, all connected to each other, and secondly, have positive edge weights. We use a standard method in graph theory – the Bron-Kerbosch algorithm [21] – to find sets of AUs that satisfy these two conditions. We consider any two AUs, *i* and *j* to be connected with a positive edge weight if both 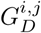 and 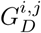 have a positive value. For every set we sum its positive edge weights in *G*_*D*_ and select the set with the highest sum as our final conjoined pain AUs.

### 4.3 Observation Window Size (OWS)

The Observation Window Size determines how close in time two AUs must occur to be considered as co-occurring. For example if two AUs occur within the same 5 second slice, with a *OWS* = 5, they would be counted as co-occurring. With longer OWS, more AUs will co-occur.

Our total dataset comprises of twelve 30 second long video clips. We use a sliding window based approach to split each video into shorter clips where the step size is set to half the OWS. For example with *OWS* = 5 a 30 second video will be split into 11 shorter clips of duration 5 seconds, starting at times 0, 2.5, 5, 7.5 and so on, seconds. We explore OWS set to 2, 5, 10, 15, 20 and 30 seconds.

By exploring OWS of increasing length, we can capture AU co-occurrence dynamics of varied time length. Each of these shorter clips is treated as a separate pain or no-pain observation. A smaller OWS helps increase the size of our dataset so that more reliable assertions can be made.

## 5 Results

### 5.1 Human FACS Interpretation (HFI)

Table 1 summarizes the AUs that passed the frequency and distinctiveness criterion for selection, along with the percentage of total AU occurrences each comprised, and the percentage difference in frequency each exhibited between pain and no-pain videos.

**Table 1.** AUs found to be associated with pain using the Human FACS Interpretation Method.

| Action Unit | AU17<br>(Chin Raiser) | AD38<br>(Nostril Dilator) | AU47<br>(Half Blink) | EAD104<br>(Ear Rotator) | AD1<br>(Eye White Increase) | AU101<br>(Inner Brow Raiser) | AU145<br>(Blink) | EAD101<br>(Ears Forward) |
| --- | --- | --- | --- | --- | --- | --- | --- | --- |
| Percentage of all Pain video AUs | 7.23% | 10.54% | 12.35% | 13.86% | 5.72% | 13.25% | 7.83% | 8.73% |
| More Frequent in Pain Videos | ✓ | ✓ | ✓ | ✓ | ✓ | ✓ | ✗ | ✗ |
| Percentage Difference | 90.91% | 69.23% | 56.25% | 42.11% | 17.14% | 2.30% | -14.29% | -18.75% |

*Inner brow raiser* (AU101), *half blink* (AU47), *chin raiser* (AU17), *ear rotator* (EAD104), *eye white increase* (AD1), and *nostril dilator* (AD38) were associated with pain, while, of the 5% most frequent action units/descriptors, *blink* (AU145) and *ears forward* (EAD101) were not. Of the selected AUs the most pronounced percentage difference in pain and no-pain frequency is for AU17 at 90.91%, while AU101 was barely more frequent in pain videos at just 2.3%.

### 5.2 Co-Occurrence Method

Unlike the HFI Method, the Co-Occurrence method for feature selection relies on temporal information to determine pain AUs. For each OWS we determined the relevant AUs and also reported their p-value. Table 2 shows the AUs selected for each observation window size.

**Table 2.**
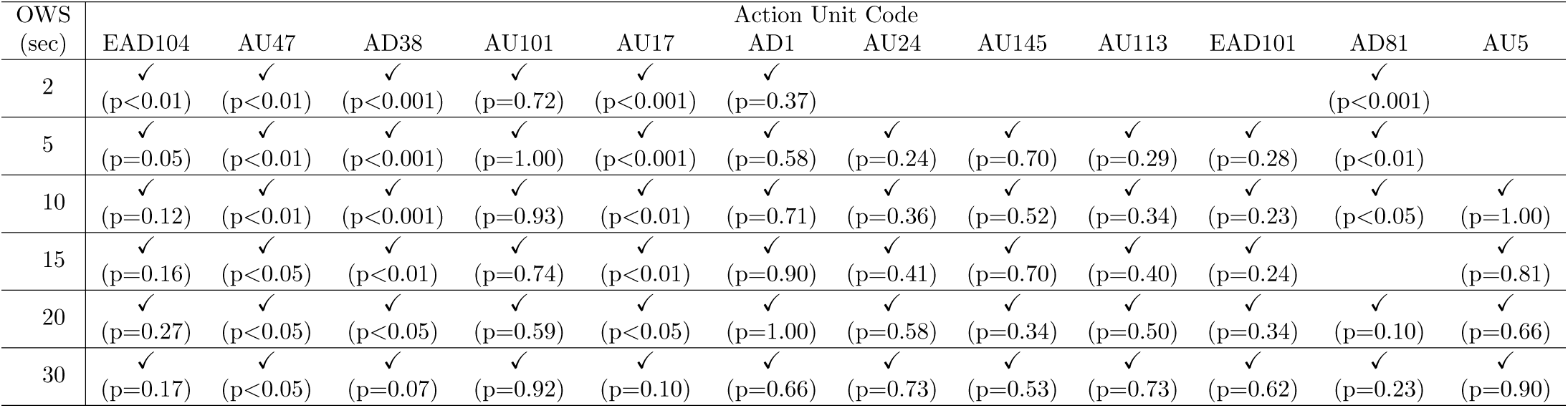
AUs selected by the Co-Occurrence method for pain detection for each Observation Window Size. Values in paranthesis show p-value using paired t-test for mean values.

*Eye white increase* (AD1), *chin raiser* (AU17), *nostril dilator* (AD38), *half blink* (AU47), *inner brow raiser* (AU101), and *ear rotator* (EAD104) are selected across all observation window sizes. All of the selected AUs are selected across multiple observation window sizes.

Of the AUs chosen across all OWS, *half blink* (AU47), *nostril dilator* (AD38), and *chin raiser* (AU17) are statistically significant – i.e. with *p <* 0.05 – across almost all OWS. On the other hand, *inner brow raiser* (AU101), and *eye white increase* (AD1) fail to show statistical significance across any observation window size. This is echoed in findings from Section 5.1, where *inner brow raiser* (AU101) is barely more frequent in pain videos compared to no-pain videos, and *eye white increase* (AD1) barely constitutes more than 5% of AU occurrences in pain videos.

Using a smaller observation window size not only accounts for briefer periods of pain expression, but also increases the number of data points for analysis. As a result, ∼ 71% of AUs selected with an OWS of 2 seconds show statistical significance. In contrast only one, or ∼ 8%, of selected AUs show statistical significance when using an observation window size of 30 seconds.

*Chewing* (AD81), demonstrates statistical significance, and is chosen as a pain AU across almost all OWS. AD81 is not a frequent action unit, constituting just 2.11% of AU occurrences in pain videos. However, its inclusion demonstrates that it occurs together with other pain AUs and is therefore important.

### 5.3 Conjoined Pain AUs

As described in Section 4.2.1, the conjoined pain AUs occur together in the same time slice, and as a group are more frequent in pain rather than no-pain instances. For brevity, we provide results for *OWS* = 2 seconds.

*Nostril dilator* (AD38), *chewing* (AD81), *upper lip raiser* (AU10), *chin raiser* (AU17), and *lip pucker* (AU18) are selected. The method selects both the AUs that demonstrate the strongest association to pain with both the HFI and Co-Occurrence methods – *nostril dilator* (AD38), and *chin raiser* (AU17). Additionally, AUs associated with lower face movement – *lip pucker* (AU18), *chewing* (AD81) – and nostril movement – *upper lip raiser* (AU10) – are selected. This indicates that lower mouth movement and nostril dilation are strong indicators of pain.

### 5.4 Specific AUs

As discussed in Section 5.1, *inner brow raiser* (AU101) is only slightly more frequent in pain videos than in no-pain videos, with a percentage difference of 2.3%. This may be due to the small size of our experimental pain dataset. For the clinical dataset, AU101 has a much higher percentage difference of 15.38%. This large difference may be due to increased levels of stress in clinical horses, which was not the case for experimental horses.

*Chin raiser* (AU17) and *nostril dilator* (AD38) are selected as AUs indicative of pain by all methods described. As a simple test, we use their presence as an indicator of pain and evaluate performance on clinical data.

Table 3 (top) shows the positive predictive value (PPV) and negative predictive value (NPV) for pain prediction for each observation. In addition, we report video level results, where the pain prediction of the of majority observation windows determines the pain prediction of the entire video. In either case, the presence of both AU17 and AD38 has a high positive predictive value for all *OWS <* 20. In particular, observing both AUs within the same 15 second interval has an 80% chance of correctly identifying pain. If majority of 15 second intervals in a 30 second interval show AU17 and AD38 co-occuring, then there is a 100% chance of the observation belonging to a pain episode. On the other hand, the absence of both AU17 and AD38 is also a fairly good indicator of no-pain, particularly for *OWS >* 5. Around 7 out of 10 observations where both AUs are absent correctly correspond with no-pain. However around 3 out of 10 times, a pain observation is incorrectly labeled as no-pain. Outside of a clinical setting, when observed horses are expected to be healthy, the absence of both AU17 and AD38 can be useful to correctly rule out the occurrence of pain. Within a clinical setting, the presence of both AUs can be used as a very specific indicator for the presence of pain.

**Table 3.** Positive and negative predictive value for different OWS on clinical data. The criteria for determining pain is the presence of AU 17, AD 38, either, or both. Missing values indicate no observation with required criteria was present.

| OWS | Positive Predictive Value (PPV) |  |  |  | Negative Predictive Value (NPV) |  |  |  |
| --- | --- | --- | --- | --- | --- | --- | --- | --- |
|  | AD38 | AU17 | Either | Both | AD38 | AU17 | Either | Both |
| 2 | 38.10% | 61.54% | 39.11% | 85.71% | 70.03% | 67.92% | 71.30% | 67.28% |
| 5 | 35.71% | 54.55% | 35.97% | 77.78% | 69.52% | 68.90% | 70.65% | 68.47% |
| 10 | 33.82% | 53.85% | 33.33% | 83.33% | 67.57% | 69.57% | 66.67% | 69.70% |
| 15 | 32.56% | 50.00% | 31.25% | 80.00% | 65.00% | 69.81% | 60.00% | 70.69% |
| 20 | 32.26% | 50.00% | 30.30% | 66.67% | 63.64% | 70.59% | 55.56% | 72.22% |
| 30 | 31.25% | 40.00% | 27.78% | 66.67% | 60.00% | 68.75% | 33.33% | 72.22% |

| OWS | Positive Predictive Value (PPV) |  |  |  | Negative Predictive Value (NPV) |  |  |  |
| --- | --- | --- | --- | --- | --- | --- | --- | --- |
|  | AD38 | AU17 | Either | Both | AD38 | AU17 | Either | Both |
| 2 | 37.50% | - | 44.44% | - | 69.23% | 66.67% | 75.00% | 66.67% |
| 5 | 30.77% | 100.00% | 33.33% | - | 62.50% | 70.00% | 66.67% | 66.67% |
| 10 | 35.71% | 50.00% | 33.33% | 100.00% | 71.43% | 68.42% | 66.67% | 70.00% |
| 15 | 33.33% | 50.00% | 31.25% | 100.00% | 66.67% | 70.59% | 60.00% | 73.68% |
| 20 | 31.25% | 40.00% | 27.78% | 66.67% | 60.00% | 68.75% | 33.33% | 72.22% |
| 30 | 31.25% | 40.00% | 27.78% | 66.67% | 60.00% | 68.75% | 33.33% | 72.22% |

### 5.5 Probability of Observing Pain

We record the percentage of observations of fixed time length where a given number of AUs associated with pain are found. We use *chin raiser* (AU17), *nostril dilator* (AD38), *half blink* (AU47), *inner brow raiser* (AU101), *eye white increase* (AD1), and *ear rotator* (EAD104) as our pain AUs since they are selected by both the Co-Occurrence, and HFI methods. In addition to the observation window sizes considered previously, we also use an observation window size of 0.04 seconds as a proxy for still image based observation since it corresponds to a frame in a 25 frames per second film. Results for pain and no-pain videos for experimental data are shown in Table 4.

**Table 4.** Percentage of observation windows from experimental data with increasing number of pain AUs present.

| Experimental No Pain Videos |  |  |  |  |  |  |  |
| --- | --- | --- | --- | --- | --- | --- | --- |
| Number of AUs | Observation Window Size (Seconds) |  |  |  |  |  |  |
|  | 0.04 | 2 | 5 | 10 | 15 | 20 | 30 |
| $\geq 1$ | 74.07% | 91.95% | 98.48% | 100.00% | 100.00% | 100.00% | 100.00% |
| $\geq 2$ | 17.58% | 65.52% | 84.85% | 96.67% | 100.00% | 100.00% | 100.00% |
| $\geq 3$ | 1.31% | 26.44% | 62.12% | 83.33% | 94.44% | 91.67% | 100.00% |
| $\geq 4$ | 0.27% | 11.49% | 30.30% | 60.00% | 66.67% | 75.00% | 83.33% |
| $\geq 5$ | 0.00% | 4.02% | 15.15% | 30.00% | 50.00% | 58.33% | 66.67% |
| 6 | 0.00% | 1.15% | 3.03% | 10.00% | 27.78% | 41.67% | 50.00% |

| Experimental Pain Videos |  |  |  |  |  |  |  |
| --- | --- | --- | --- | --- | --- | --- | --- |
| Number of AUs | Observation Window Size (Seconds) |  |  |  |  |  |  |
|  | 0.04 | 2 | 5 | 10 | 15 | 20 | 30 |
| $\geq 1$ | 81.67% | 97.70% | 100.00% | 100.00% | 100.00% | 100.00% | 100.00% |
| $\geq 2$ | 31.93% | 81.03% | 96.97% | 100.00% | 100.00% | 100.00% | 100.00% |
| $\geq 3$ | 6.13% | 59.20% | 84.85% | 96.67% | 100.00% | 100.00% | 100.00% |
| $\geq 4$ | 0.31% | 28.16% | 72.73% | 90.00% | 100.00% | 100.00% | 100.00% |
| $\geq 5$ | 0.00% | 4.02% | 24.24% | 60.00% | 66.67% | 66.67% | 66.67% |
| 6 | 0.00% | 1.15% | 7.58% | 23.33% | 44.44% | 66.67% | 66.67% |

The probability of capturing multiple pain features in a still image is very low, and longer observation time periods – video – is necessary to determine the presence of pain expression. The likelihood of observing at least 3 pain AUs is negligible in still frames at ∼ 6%, and less than a hundredth chance of observing 4 or more pain AUs. Furthermore, using still images removes observers’ ability to observe facial *changes*, so that facial behaviors that are constantly present are erroneously considered important, and minor facial changes that are important are ignored. Examples are *chin raiser* (AU17) and *half blink* (AU47), that are found to be important for pain by both the Co-Occurrence and HFI methods, but are virtually impossible to observe in a still image.

The likelihood of observing a range of pain AUs is not negligible in no-pain videos. For example, while 60% of 10 second pain clips display 5 or more pain AUs, 30% of no-pain 10 second clips also display 5 or more pain AUs. This implies that reliable pain detection may firstly, require consolidating information from multiple observations, and secondly, require a more sophisticated method for evaluating pain than summing the number of observed pain features. For instance, AU17 is only present in ∼ 16.7% no-pain observation windows of 10 second length, but 50% of 10 second pain segments. Weighting AUs differently may hence be beneficial for reliable pain detection.

## 6 Discussion

The dataset produced by the fine-grained method of EquiFACS describes the temporal emergence and disappearance of 17 facial action units and 26 action descriptors over 30 seconds of video (see S13). However, data driven methods do not exist for the interpretation of such data in horses or other animals. We therefore explored statistical methods used in human FACS [17] and also developed novel statistical approaches to analyze the FACS data.

This study describes for the first time the facial activities of horse in pain by use of EquiFACS. Using the HFI method, the characteristic AUs for horses in pain were *chin raiser* (AU17), *nostril dilator* (AD38), *half blink* (AU47), *inner brow raiser* (AU101), *eye white increase* (AD1), and *ear rotator* (EAD104). An image of a horse head showing these features, excluding the “half blink” (AU47) which is not possible to illustrate in an image, would probably be assessed as a “Pain Face” according to the description in [11], and it would achieve scores indicative of “pain” using the Horse Grimace Scale [10] and the FAP scale [8], showing concurrent validity of our results, when compared to clinical empirical methods.

The two actions most prevalent in painful horse were the *chin raiser* (AU17) and *nostril dilator* (AD38). These two facial actions seem to have equivalents in the Horse Grimace Scale [10] as the configurations “mouth strained and pronounced chin” and “strained nostrils and flattening of the profile”; in the Facial Assessment of Pain scale [8] as the configuration (nostrils) “A bit more opened” or “Obviously more opened, nostril flaring” and “Corners mouth/ Lifted a bit” or “Obviously lifted”, and in the Pain Face as the configuration “Edged shape of the muzzle with lips pressed together” and “Nostril dilated in the medio-lateral direction”. This indicates a concurrent validity of our results, when compared to clinical empirical methods.

The third most prevalent action of the face was the *half blink* (AU47), which is defined as a reduction of the eye opening by the eyelids, but without complete closure of the eye [14]. This blink has not been described as an indication of pain before, probably because it is only possible to appreciate this feature from close inspection of video. The action takes place in less than half a second [14]. Decreased eye blink rates has recently been associated to occurrence of stress or pain [22]. The Horse Grimace Scale [10] describes “Orbital Tightening: The eyelid is partially or completely closed”. It is not clear whether this corresponds to an “eye closure” (AU 45) with a duration of more than half a second or if the rapid half blink is part of this definition. In opposition to this, the FAP scales [8] uses closure as the eye and opening of the eye as an indicator for pain, and relates intensity of pain to the amount of sclera that can be seen. While the Pain Face illustration can not describe blinks, half blinks or eye closure, the ethogram behind it contains clear evidence of increased blinking behaviour as indicative of horses in pain [11]. The *inner brow raiser* (AU101) is also associated to characteristic changes in the eye region. The *inner brow raiser* is said to create a “triangular eye” or “worry wrinkles” [22, 23], an appearance traditionally related to both stress and pain by horse care providers and veterinarians. Per definition, this muscle contraction increases the perceived size of the eye region, but not the aperture of the eye [14]. This activity is also used in the Horse Grimace Scale and described as “tension above the eye area” [10], and as “contraction of m. levator anguli oculi medialis” in the Equine Pain Face [24]. It is therefore surprising that AU101 was only barely more often shown in the pain group of this study. The acclimatization of the horses in the present study may have led to a generally lower level of stress, indicating that AU101 may be a sign of general stress and pain, since these conditions seem inseparable.

In the FAP scale, a configuration around the eye is described as “obviously more opened eyes”, and also increased visibility of the sclera, the “eye white” [8]. In EquiFACS these features would be coded as *upper lid raiser* (AU5), and *eye white increase* (AD1), the latter being associated to pain, though it did not achieve statistical significance.

The ears are highly communicative in horses [25] and these FACS codes naturally have no human parallel. In this study, *ear rotator* (EAD104) was associated with pain. In the Horse Grimace Scale a “moderately present-stiffly backwards ear” resembles EAD 104, and the “obviously present stiffly backwards ear” with a wider distance between the tips of the ears resemble the *ear flattener* (EAD103), which has another muscular basis [10]. In the description of the Pain Face, “the lowered ears” with a broader base resembles the EAD 104, while the “asymmetric ears” described in the Pain Face have no single equivalent in EquiFACS [24]. The FAP scale uses the “backwards ears” – it is not clear if EAD 104 or EAD 103 are parallels, or both [8].

In summary, the EquiFACS and the simple frequency methods applied from human research seem to point out a number of facial configurations already described in other pain studies. One important difference was the occurrence of the *half blink* (AU47), which has never specifically pointed out as a facial expression of pain. An explanation for this may be that this feature is extremely difficult to appreciate by life scoring or from images. Also important were discrepancies regarding the role of the configurations related to the eye.

The decision for diagnosing pain in horses by human observers are typically based on prototypical configurations (grimaces), where some can be atomic movements like facial action units, while other are more broadly described. We therefore investigated other methods that would take into consideration the co-occurrence of the facial action units and not only their frequencies.

The Co-occurrence method is a novel method, where we used observation windows of varying time lengths to investigate which facial actions occurred together in any given range of time in each video sequence.

A more complex picture emerged when the Co-occurrence method was applied. Many more action units were selected, as shown in Table 2. In particular, *chewing* (AD81) was found to be important, despite low frequency. *Eye white increase* (AD1), and *inner brow raiser* (AU101), although selected across all observation time lengths, showed low statistical significance. The importance of mouth and nostril movement in pain detection was further highlighted when using the Co-Occurrence graphs to determine “Conjoined Pain AUs”. Here, high frequency of occurrence was not an implicit or explicit criteria for AU selection. In addition to the *chin raiser* (AU17), *nostril dilator* (AD38), and *chewing action* (AD81) identified previously, the *lip pucker* (AU18), and *upper lip raiser* (AU10) were also identified as indicative of pain.

It is also be speculated if some of these action units are associated to pain intensity. While the clinical scales contain intensity scores, as do the human FACS system [13] this has yet to be developed for the EquiFACS.

During clinical conditions, it may be important to exclude the presence of pain in a horse. In this respect, the absence of *nostril dilator* (AD38) and *chin raiser* (AU17) may be used as a proxy for no-pain if expectation of pain is low, and their presence may be used as a very reliable proxy for pain (high positive predictive value).

When clinical pain video were analyzed, the difference between occurrence of *inner brow raiser* (AU101) was much larger between pain and no-pain horses than in the experimental setting. This may be due to the overlay of stress and pain in clinical settings, that was absent from experimental data. Interestingly, the positive predictive values of *nostril dilator* (AU38) and *chin raiser* (AU 17) were perfect if the actions were seen together within a range of 10 to 15 seconds. The absence of these actions had a poorer negative predictive value, meaning that other actions should be looked for if a horse should be claimed without pain.

Important temporal information has been shown to be lost if frames are randomly scored [26]. We therefore recommend that facial actions are scored from video, with onset and offset noted, by trained and certified raters. We were surprised to discover that if a pain assessment was depending on only one randomly picked frame (0.04 seconds duration) from the pain videos, the chance that more than three actions units were present would be ∼ 0.3%, meaning that there was high risk that the painful horse would not be assessed as in pain. On the other hand, if one frame were picked from the no-pain group of horse, the chance of assessing this erroneously as a horse in pain would be quite similar at 0.27%. The random picking of frames from videos therefore has very poor predictive values, and can not be recommended as a method. Selection of images and pain assessment from photographs showing “typical features” thus holds a high risk of providing biased and thus wrong conclusions. Additionally, some AUs, such as *half blink* (AU47), cannot be reliably scored from still frames.

The limitations of this study is the low number of horses and the experimental conditions, where horses were acclimatized to external inputs. Further, this study only investigated the facial activities produced by a single pain modality. Despite the fact that facial expressions have been shown to be one of the best suitable pain indicators in people with dementia, who also can not self-report [27], pain should be studied in more horses, during clinical conditions and with different types of pain.

In conclusion we have for the first time described the facial activities of one “prototypical” pain face of acute pain in the horse using an EquiFACS protocol and both traditional and new statistical methods. We defined the *half blink* as a new indicator for pain in horses, and raised some doubts about the pain indicating value of the *inner brow raiser*. Such findings may be used to refine and further validate current horse pain evaluation scales.

## Acknowledgments

DVM Johan Lundblad, and DVM Camilla Frisk are thanked for expert rating videos of horses for pain.

## 7 Supplementary Materials

**S1-12** : Experimental Pain Data Horse Videos 1-12.

**S13** : Experimental Pain Data FACS, Pain, and Horse ID annotation for Videos 1-12.

**S14** : Clinical Pain Data FACS, and Pain annotation for 21 clinical videos.

